# Depth Dataset Using Microsoft Kinect-v2

**DOI:** 10.1101/2021.02.04.429850

**Authors:** Hamed Heravi, Masumeh Delgarmi, Ali Rahimpour Jounghani, Abdollah shahrezaie, Afshin Ebrahimi, Mousa Shamsi

## Abstract

In biomedical imaging studies, numerous methods have been used to capture human data, mostly by using magnetic resonance imaging (MRI) and computed tomography (CT). However, due to being inexpensive and accessibility of Microsoft Kinect, its usage has been significantly increased in recent years. In this study, we aimed to represent the procedure of data acquisition, which includes a set of depth images from individuals’ back surfaces. The goal of image acquisition is to investigate spinal deformities and landmark detection of the back surface. Traditional imaging systems are challenging, most notably because of ionized beams in the data acquisition process, which has not been solved yet. Indeed, noninvasiveness is the most crucial advantage of our study. Our imaging system was set in a dim laboratory, and the University approved the ethical letter of Medical Sciences before data acquisition. After that, the subjects (total 105; 50 women and 55 men) were recruited, and data images were captured from the back surface. Then, we increased the imaging data size by using the augmentation method to use deep learning methods in future works. Finally, this Dataset leads us to the desired output in our study procedure.

## 1 Introduction

Since the advent of Microsoft’s Kinect sensor, many databases with depth images have been published. Recent progress in this sensor has provided many opportunities in various fields [1]. This sensor was initially made to control video games, but later, it became a powerful and well-known tool for scientific research to achieve high-quality depth data [2] quickly. Hence, due to its low price, quick access, ease of use, and decent quality, Kinect has attracted researchers’ attention. In recent years, most of the databases derived from this sensor have been made available in various fields, including computer graphics, image processing, machine vision, object recognition, surface tracking, modeling, and robotic vision. It is of high importance for comparing and testing the most up-to-date methods [3]. The advantages of depth images include resistance to changes in color, brightness, rotation angle, and scale [4]. This research’s main objective is to collect appropriate depth images of human back anatomy using the Kinect-v2 camera for medical applications, including detection of anatomical landmarks of the human back surface, followed by the diagnosis of spinal deformities. The database contains 210 depth images taken from 105 people with fixed gestures. In general, this paper aims to provide a comprehensive description of this database. In Section 2, we will review the Kinect sensor and evaluate its hardware specifications. In Section 3, we will explain the registration and imaging stages, and, in the final section, we will focus on Groundtruth generation to use this database to detect anatomical landmarks.

### 2 A brief review of Kinect

In the last decade, RGB-D cameras, also known as range-imaging cameras, have made considerable evolution. Due to their cost and capability of measuring distance at high frame rates, such sensors have received significant attention to be used in computer vision or robot vision [4]. The release of Microsoft Kinect-v1 in November 2010 improved the use of RGB-D cameras until the second version of the sensor entered the market in July 2014. Although Kinect was initially designed to be used in Xbox 360 video game consoles, its use was not limited to such applications, and this technology has been employed in other fields such as robot navigation and control, object recognition, and image processing [5].

Since access to high-frequency point clouds of the observed scene is possible, these sensors can collect 3D information. Such imaging devices utilize optical properties and are called active sensors because they use their light source to enlighten the scene [6]. However, while Kinect-v1 was based on active triangulation through structured light, the Kinect-v2 sensor was designed as a time-of-flight (ToF) camera. The nearest object’s depth size was provided for each pixel of the depth map [5,6]. In this section, Kinect will be examined from a hardware perspective.

### 2-1 Kinect hardware Configuration

Kinect-v2 sensor is composed of two cameras, namely an RGB and an infrared (IR) camera. Three IR projectors provide the illumination of the observed scene. As mentioned, this sensor is based on the ToF method; therefore, the measured distance is proportional to the time required by the active light source to move from the emitter to the target [5,6]. Some features of the sensor are presented in Table 1.

**Table 1.**
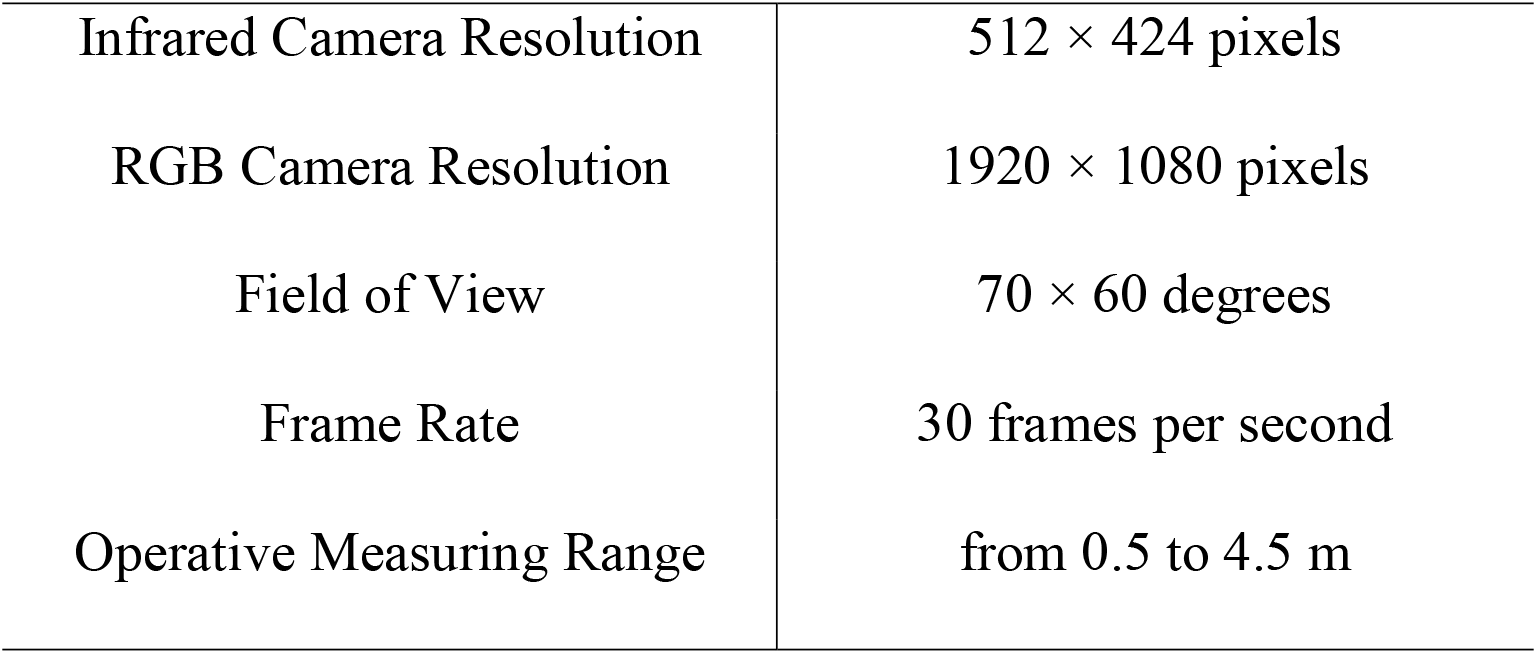
Technical features of the Kinect v2 sensor [1]

Three different outputs are obtained from two lenses of Kinect-v2 sensor: infrared data and depth maps are obtained from a single lens and have a similar resolution. Depth maps are 16-bit 2D images that store information measured for each pixel. The high-resolution color images result from the second lens [4]. Figure 1 shows a Kinect-v2 sensor with its components.

**Figure1:**
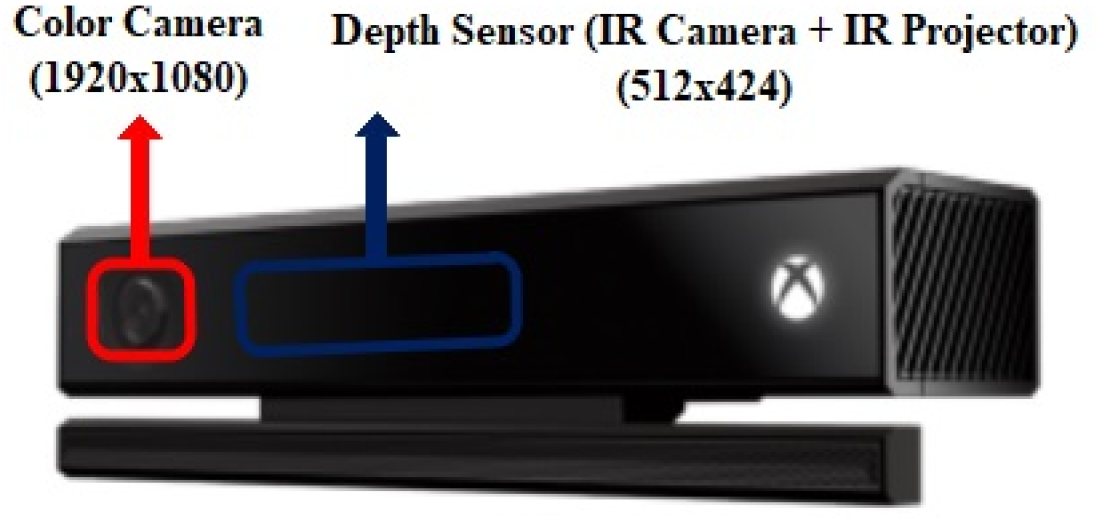
Microsoft Kinect V2

### 3 Depth Dataset

In this section, the method used to collect the depth data and create the GroundTruth from raw data will be explained.

### 3-1 Data Acquisition

The purpose of collecting this database is to detect the anatomical landmarks of the human back surface. To extract these points specified in Figure 3, an image of the human back is needed. In most of the previous studies, color images have been used to extract anatomical points in the human postural structure; however, due to the sensitivity of the problem and its use in medical applications, referring to color images as the source and input of the processing method can affect the evaluation and conclusion. Therefore, it is not appropriate to cite color images. The depth information and depth images with the capability of image registration of the human back surface and having body shape information not available in color images can report duplicate information such as ups and downs of the human back surface. This information can carry very invaluable results for diagnosis.

In this study, Microsoft’s Kinect-v2 sensor was used for data collection. Fifty women and 55 men aged 19-35 years old agreed to participate in the experiment, and the result of this collaboration was 210 depth images from the human back surface. They were asked to naturally stand with their backs to the camera with an identical distance from the feet to the plumb line. The Kinect sensor was located 1.5 m behind the participant. As shown in Figure 2, the data entry process was as follows: first, a marker was put at point C7, and imaging was done with other landmarks. After registering this image, the other markers were installed at the exact location, and the second image was registered with the presence of landmarks. The images were saved as raw images in the size of 512×424.

**Figure2:**
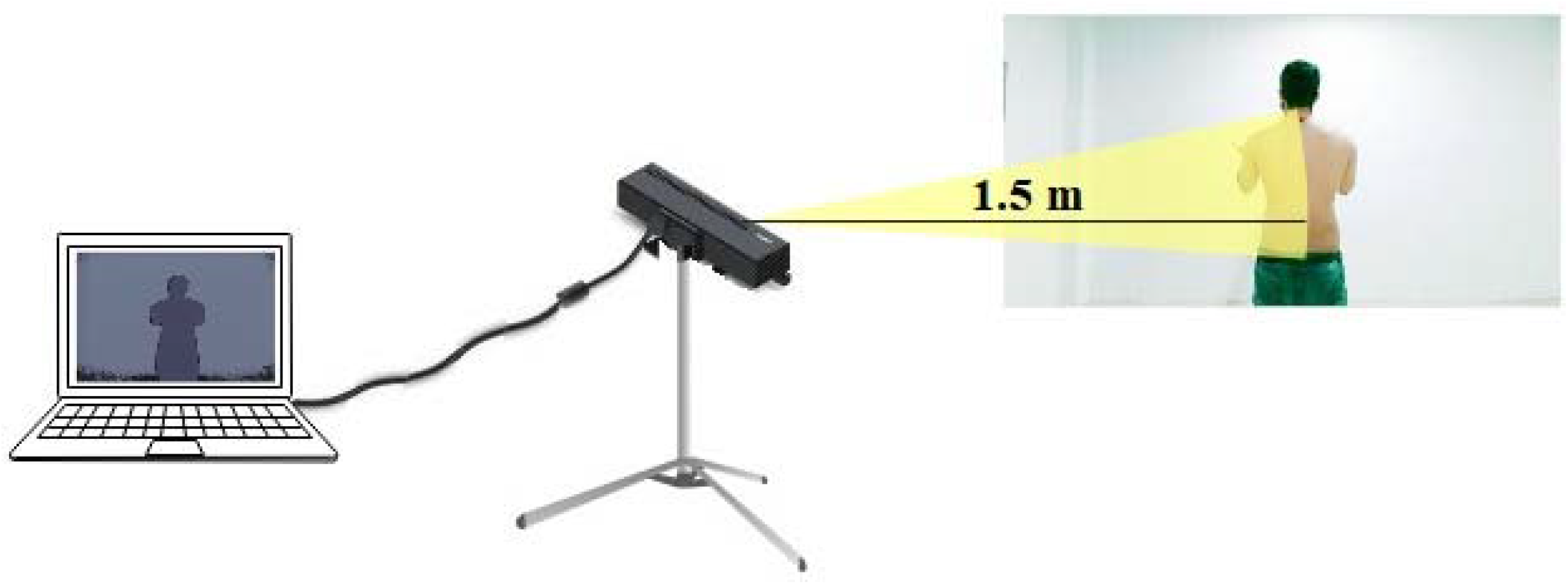
This schematic shows a human back surface captured by the Kinect sensor

The main problem with ToF cameras is that the measurements made by several phenomena are disrupted; therefore, to ensure the reliability of the obtained point clouds, this distortion must be removed in the beginning [4]. To do so, it is necessary to detect the sources of error that affect the measurement. Thus, we decided to limit the imaging conditions to a closed environment away from direct light. During image registration, the ToF camera gradually warms up, and the temperature slowly rises; the sensor’s cooling fan starts working after 20 min, and the distance from the camera remains almost fixed. Hence, before imaging, a delay of 20 min is required to keep the sensor in a stable condition [4].

### 3-1-1 Groundtruth Translation (or Annotation Technique)

GroundTruth was prepared in two steps. First step: as shown in figures 3 and 4, corresponding to each volunteer, two images have been captured: 1. Image with a marker. 2. Image without a marker (only C7 marker). Since we use images without landmarks for our study purposes, we need to access each landmark’s precise coordinates in individuals’ back surfaces using the images with markers. As depicted in figure 7, Kinect sensor output shows an unclear image of the case, therefore in this step, the color image of the subject was shown by the following structure to make the image visible. Both a2 and b2 image columns in figures 3 and 4 are the results of this strategy, indicating landmark positions.

**Figure3:**
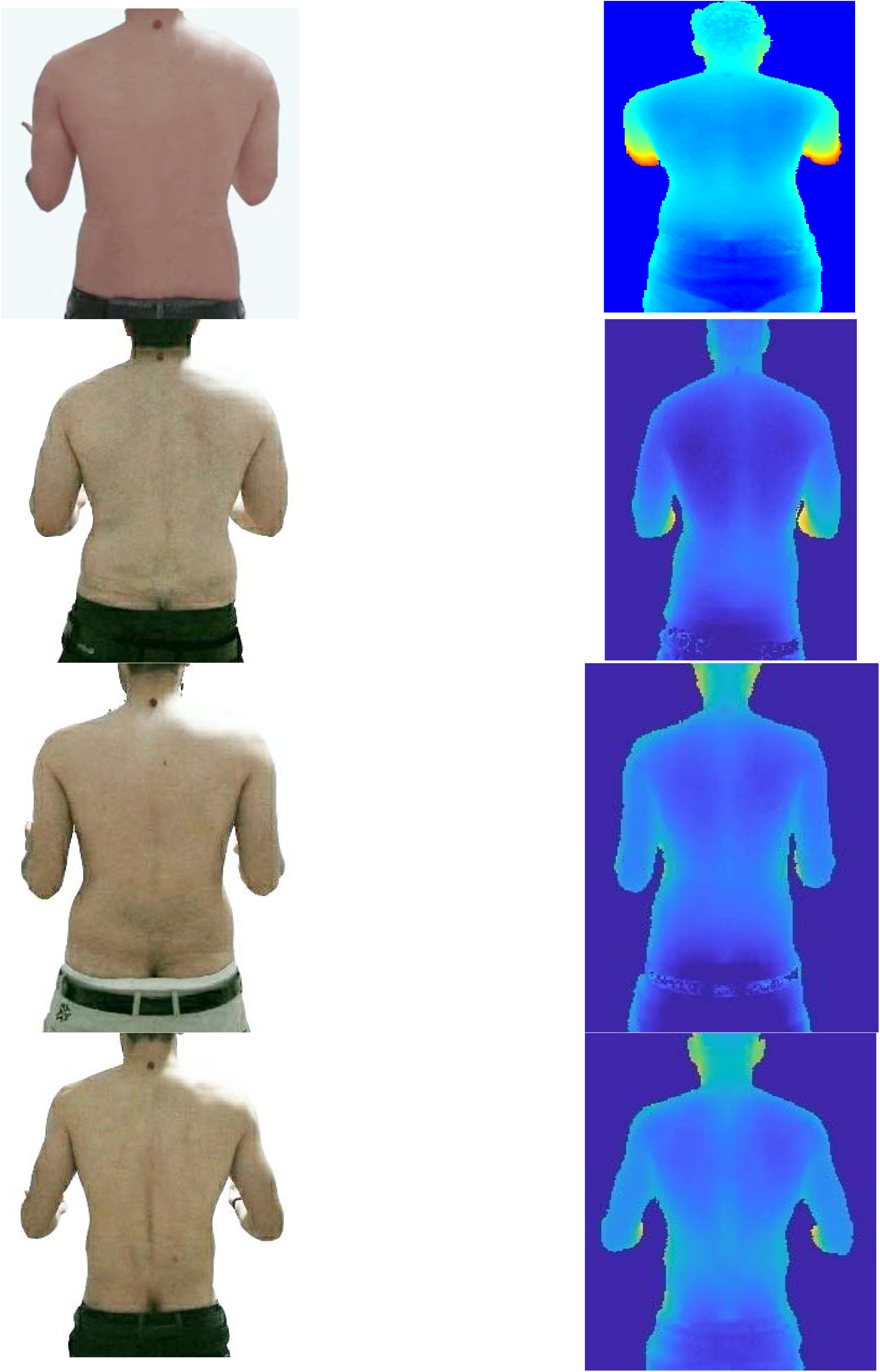
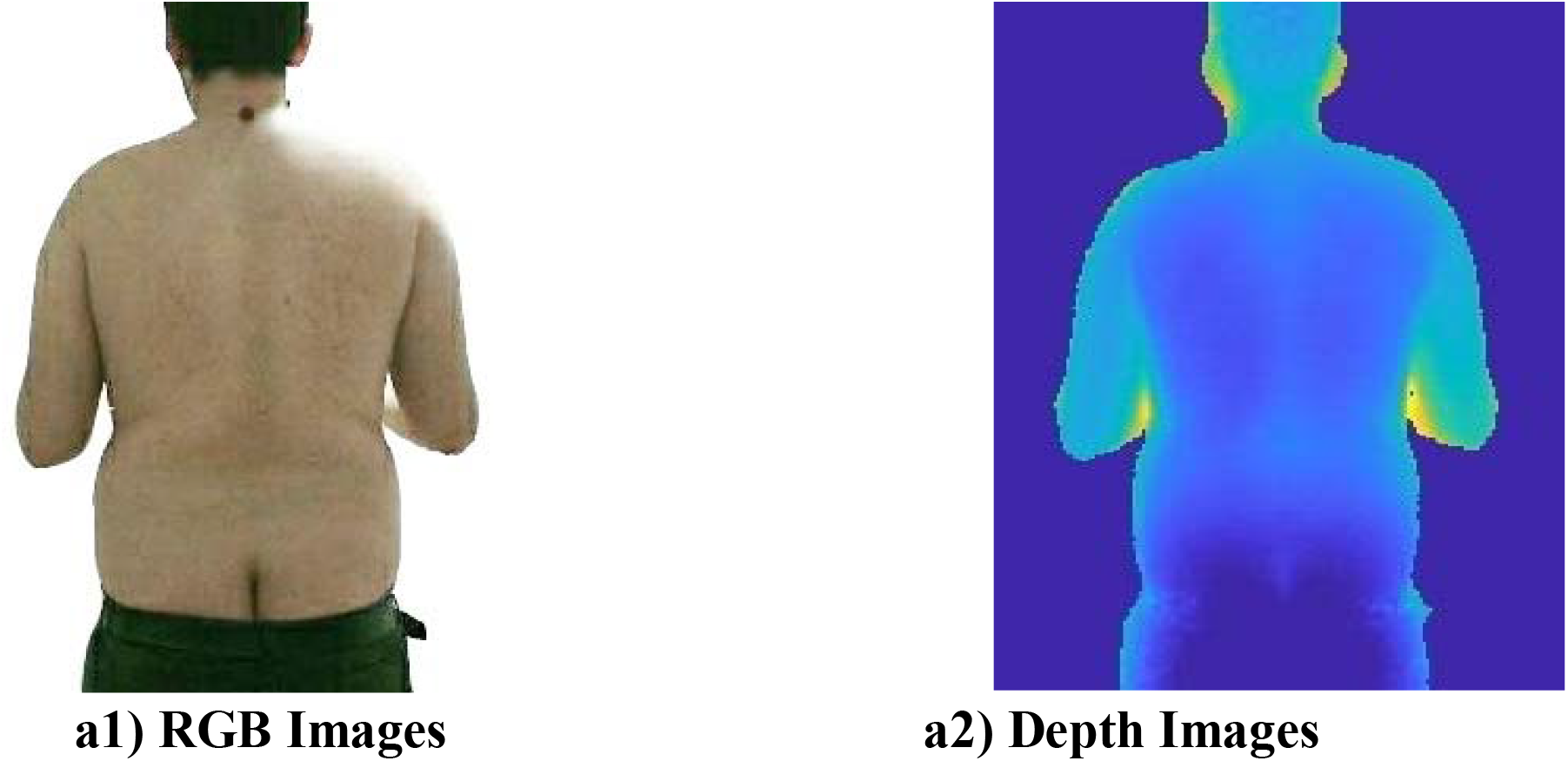
Example of captured images without markers

**Figure4:**
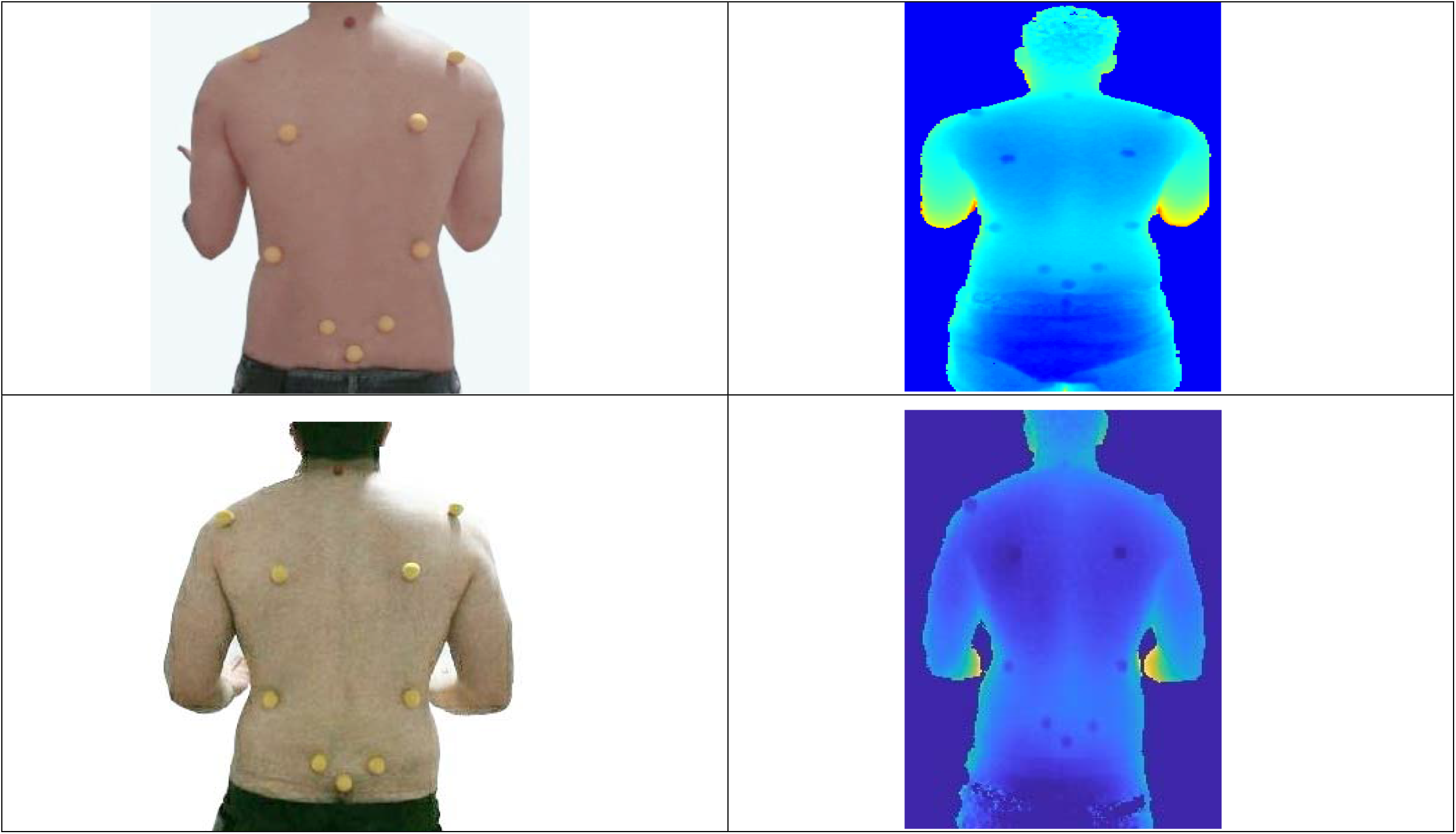
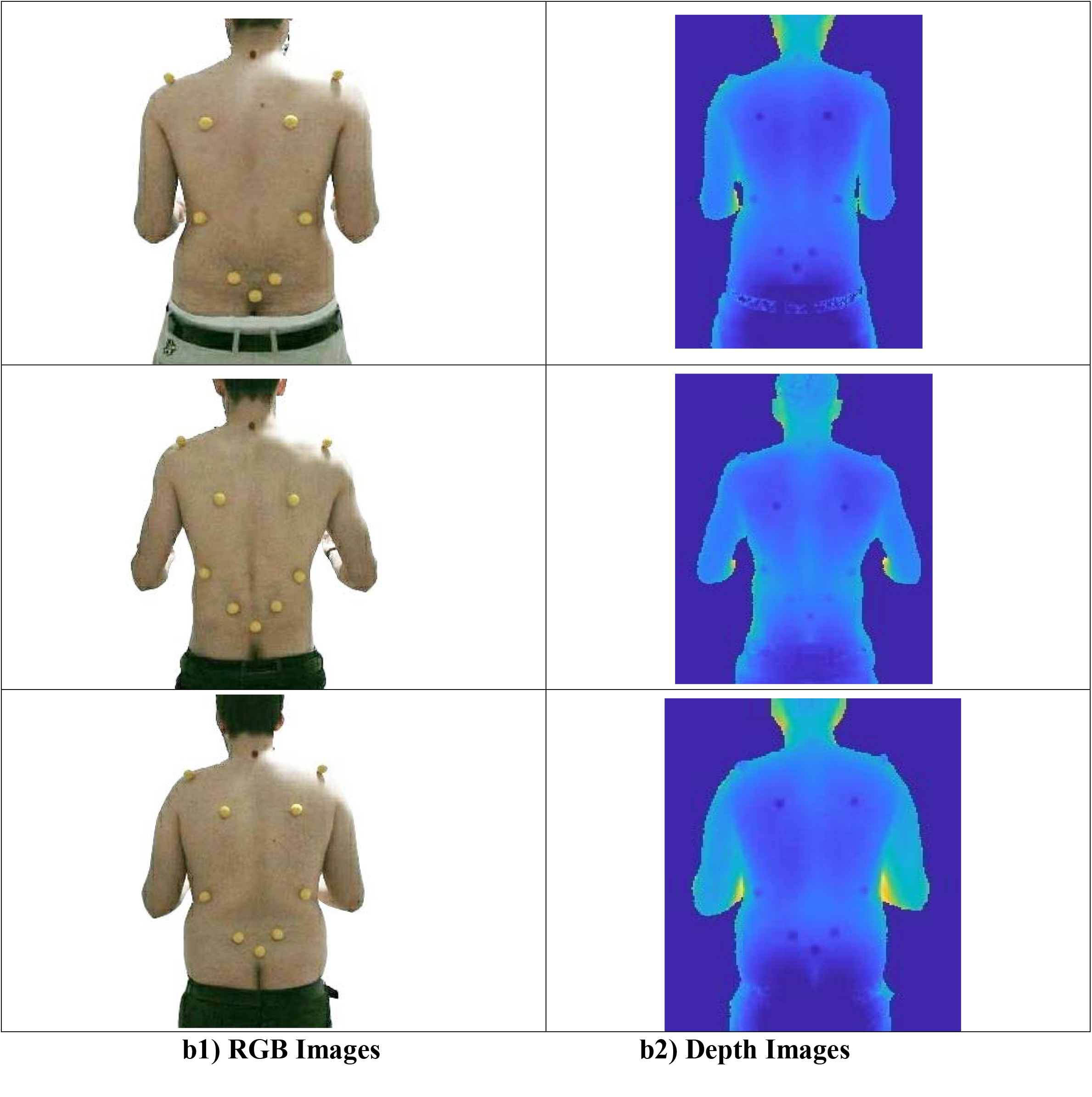
Example of captured images with markers

**Figure 5:**
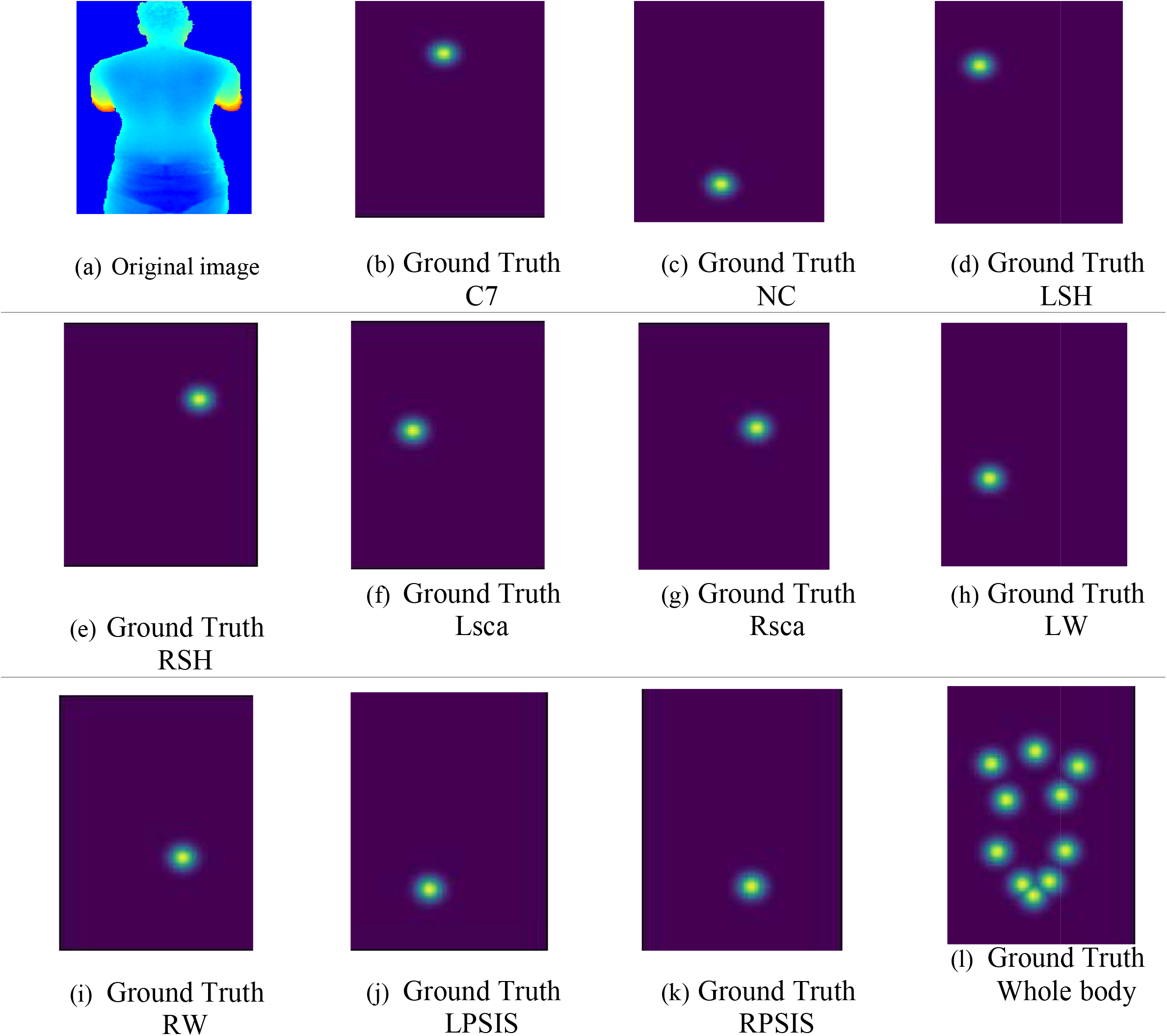
Example of Heatmaps

**Figure 6:**
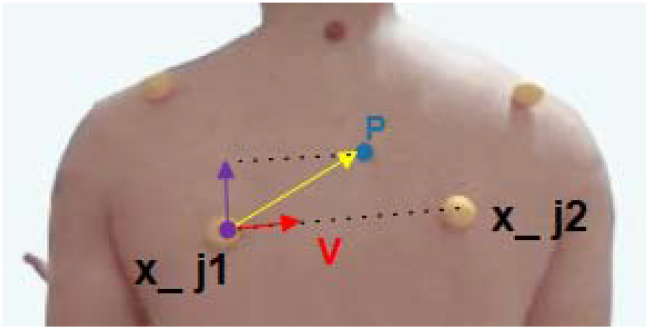
Landmark Affinity Field

Assume that *w*(*x, y*) is a pixel of the depth image. A value of each output pixel is stated in millimeters (mm). Since the camera distance with the subject is set to 1.5 (m), after trial and error, we noticed that the subjects were in the range of 1150 and 1480 (mm) on average. By using the following instruction, a visible image of the subject’s landmark was obtained.

if (w(x, y)

w(x,y)=(w(x,y)-1150)*8

*else*

Second step: In this step, we aim to create a Groundtruth to estimate landmarks’ coordinates with the Gaussian function. For each landmark, a label is generated as Groundtruth, in which the average shows the location of the individual landmarks in the image. Each heatmap indicates the pixel position in which each of the landmarks has occurred. These heatmaps are derived from the coordinates of the critical points stored in the JSON file.

Assume 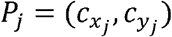 is the correct position of the j^th^ landmark for the person in the image, *x*_*j*_, *y*_*j*_ ∈ *R*^*2*^. To create a heatmap for this landmark, it is necessary to define a neighborhood to the centrality of this point because it is clear that each heatmap does not contain only one point but also covers a small area. Then, the value of the Gaussian function per *P* = (*x, y*) is calculated in this region. A heatmap for the j^th^ landmark is shown with 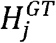. Then, the value of the Gaussian function in *P* ∈ *R*^2^ per j^th^ landmark is defined as follows [11-14]:

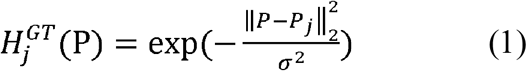

Equation 1 shows the 2D Gaussian function to the centrality of the correct landmark position. An example of the heatmaps obtained by the above method is shown in figure 1.

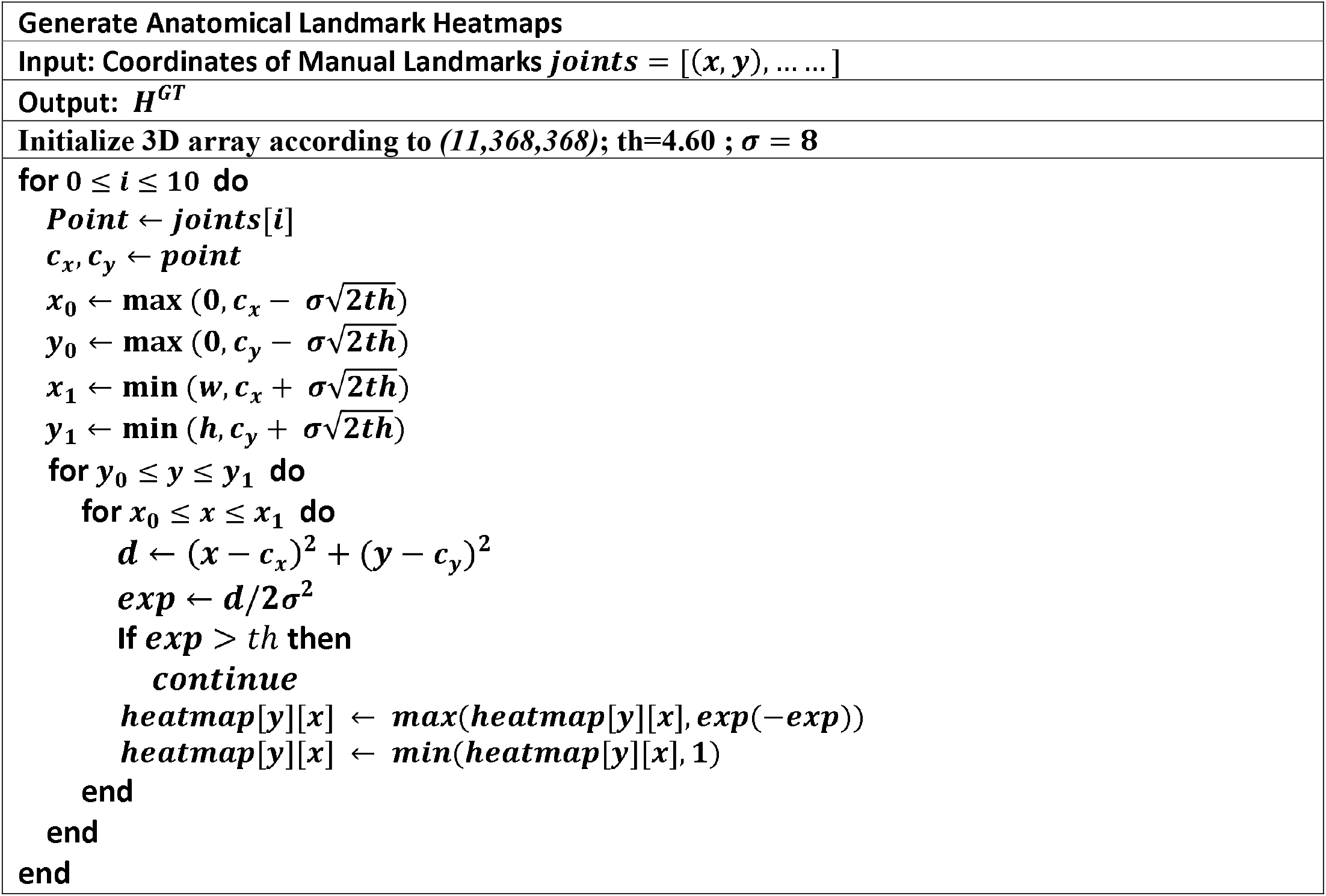

Each of the defined landmarks can be linked to the other one. Therefore, a vector map connects two parts of the body and contains position and direction information in the connection areas. In Figure 7, a connection is shown between two landmarks. Assume and are the positions of two related landmarks and in the image; if point p is on connection c, the value of is a single vector [15-17]. Otherwise, there is a vector with a value of zero for all other points [11].

A vector map at the points of image p is defined as follows [11]:

(2)

Here, v is a unit vector towards the connection and is defined as follows [11]:

__________________ (3)

Figure 7 shows an example of vector maps obtained by the above method.

**Figure 7:**
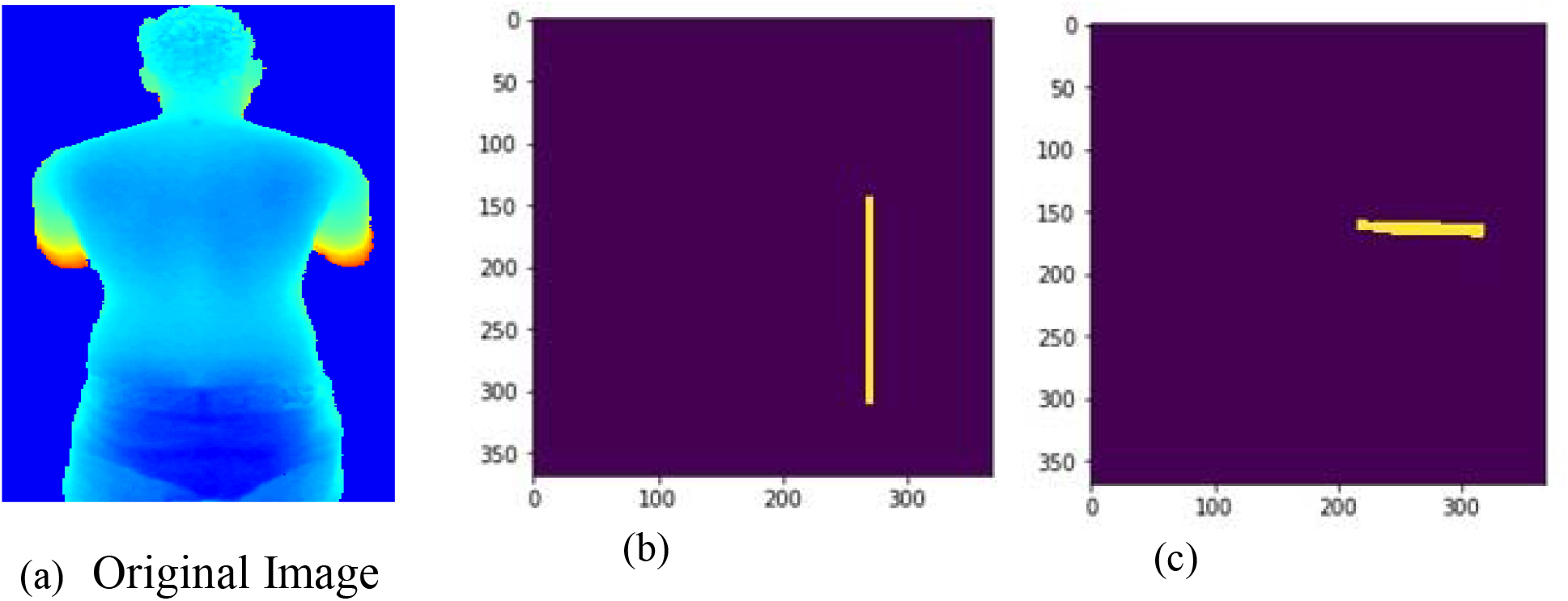
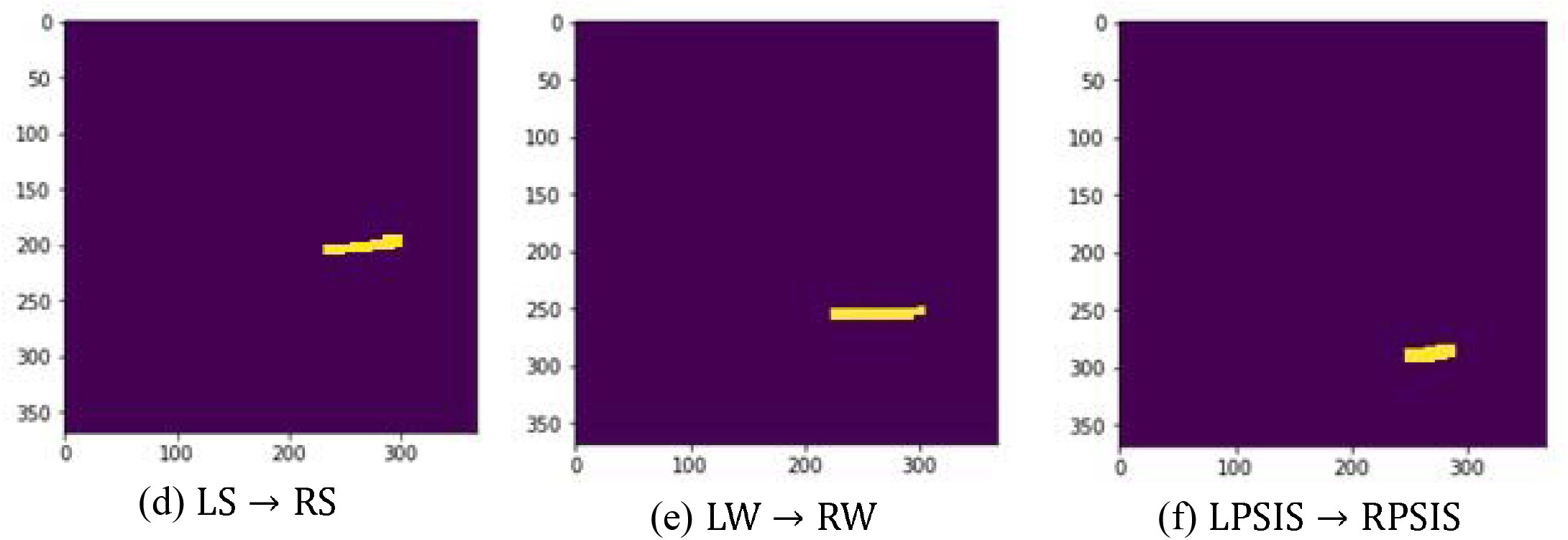
Example of vectormap

### 3-2 Denoising, Data Augmentation

Given the nature of the Kinect sensor, the derived depth images are very noisy [7]. Hence, it is necessary to apply a preprocessing step to improve the quality of the depth image. Thus, if r is the depth image matrix, *r* > 1.7*m* values are removed by thresholding. Figure 8 shows the depth image before and after noise removal.

**Figure 8:**
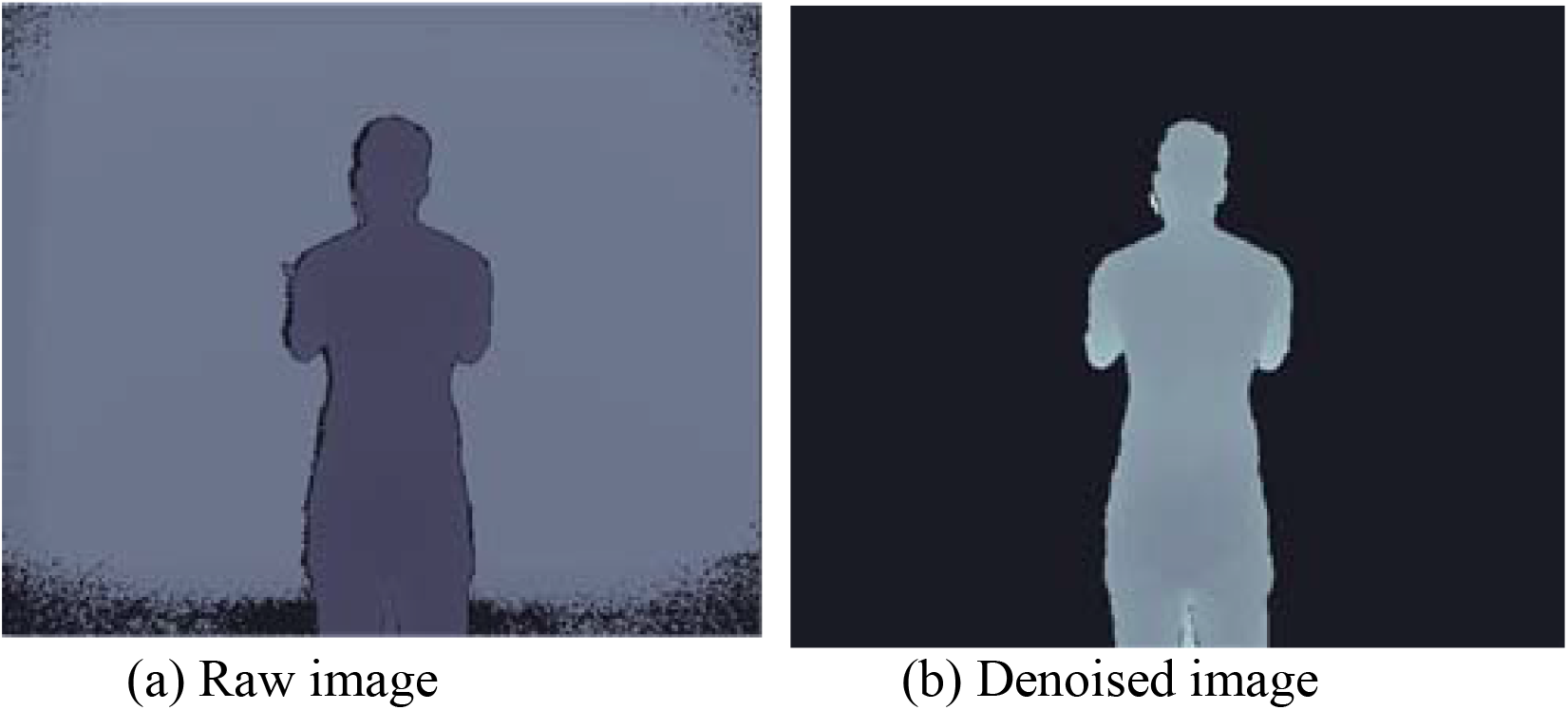
Example of Raw image and Denoised image

**Figure 9:**
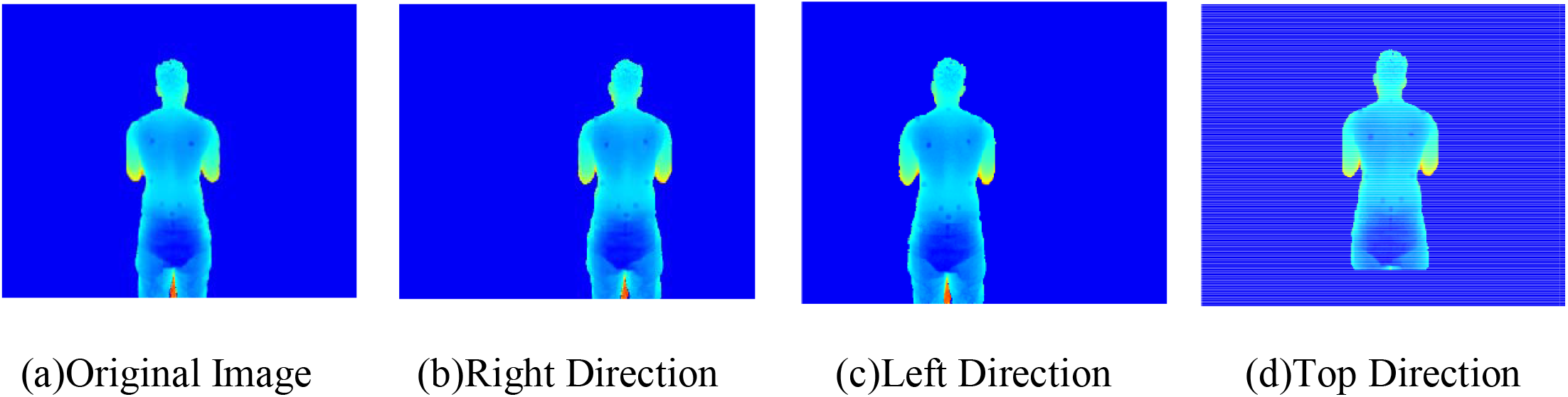
Example of Data Augmentation

If we want to use the deep learning method for specific expertise, such as detecting anatomical landmarks, there is a basic need to have an extensive database. Compared with large datasets used in deep learning, such as COCO [8], ImageNet [9], etc., the Dataset collected for medical applications that need precise detection, and high sensitivity is minimal. Then, data augmentation [6] is used to solve the problem of data shortage. We applied four transfer techniques to the right, left, top, and bottom, to the images in Dataset, and the final number of image set reached 525 images. Figure 5 shows an example of images in this database after preprocessing and data augmentation processes.

## 4 Discussion and Conclusion

In this study, we prepared a dataset named BackSurfDepth, which involves 210 depth images (105 without landmarks and 100 with landmarks). This database’s necessity is rooted in our need to access a set of coherent images for investigating spinal deformities and landmark detection of the back surface. An essential advantage of this method is using the sensors, making it non-invasive, inexpensive, and easy to access. The results from the data analysis indicated the excellence of the data for medical purposes. Therefore, we used this Dataset for automatic detection of anatomical landmarks in the back surface.

